# Dissecting cell state transitions by causal inference

**DOI:** 10.1101/2023.02.08.527606

**Authors:** Longchen Xu, TingTing Cong, Hengyi Xu, Naixue Yang, Chen Tian, Sijie Yang, Ming Zhu, Rahul Sinha, Ryo Yamamoto, Wei Zhang, Jianwei Wang, Xun Lan

## Abstract

Trajectory inference methods are frequently used for cell fate analysis, however, most of them are similarity-based and lack an understanding of the causality underlying differentiation processes. Here, we present CIBER, a <u>C</u>ausal <u>I</u>nference–<u>B</u>ased framework for the <u>E</u>valuation of feature effects and the <u>R</u>econstruction of cellular differentiation networks. CIBER provides a novel paradigm for dissecting cell state transitions other than trajectory inference and differential analysis. It is a versatile framework that can be applied to various types of data, including transcriptomic, epigenomic and microarray data. It can identify both known and potential cell-lineage structures with minimal prior knowledge. By integrating the CIBER-learned network with structural causal model and applying *in silico* perturbation as inventions, we generated an effect matrix that quantifies the impact of different features on each differentiation branch. Using this effect matrix, CIBER can identify crucial features involved in haematopoiesis, even if these features show no significant difference in expression between lineages. Moreover, CIBER can predict novel regulation associations and provide insight into the potential mechanism underlying the influence of transcription factors on biological processes. To validate CIBER’s capabilities, we conducted *in vivo* experiments on *Bcl11b*, a non-differentially expressed transcription factor identified by CIBER. Our results demonstrate that dysfunction of *Bcl11b* leads to a bias towards myeloid lineage differentiation at the expense of lymphoid lineage, which is consistent with our predictions.

## Introduction

Cellular differentiation is a fundamental process in biology that underlies the development of complex organisms. Advances in single-cell omics data have provided unprecedented opportunities for elucidating developmental pathways and reconstructing cell lineages. Computational trajectory inference methods have been developed to study cellular differentiation. In most of these methods, cells are ordered along deterministic^1–5^ or probabilistic^6,7^ trajectories based on similarities in their expression patterns, and a numeric value known as the pseudotime is assigned to each cell to indicate how far it has progressed along a dynamic process of interest. However, these similarity-based methods may not capture the causal nature of cellular differentiation. Another types of trajectory inference methods^8–12^ are RNA velocity-based, which model the differentiation process with the spliced-to-unspliced mRNA ratio in the form of differential equations, thereby restoring cell temporal information, and can then be utilized for causal gene regulatory network reconstruction^13^. However, RNA velocity need to represent cell dynamics in the transcriptome space^14^ and is not suitable for other types of data such as epigenomic or microarray data alone. Recently, new methods^15,16^ have been developed to infer how cell identity is governed by the complex regulation of gene expression through *in silico* transcription factor (TF) perturbation. However, they aim at inferencing the change in cellular differentiation process but not the process itself and need the pseudotime provided by other methods. More general approaches are required to identify causality between cellular states with minimal prior knowledge, not only through transcriptomic profiling but also through other types of data.

Granger causality^17–19^ and Bayesian network^20–23^ (BN) have been effectively applied for causal inference in studying biological processes such as gene regulatory networks and protein-protein interactions. However, these methods are rarely applied in scenarios such as the reconstruction of cellular differentiation networks and the identification of fate-determining features. BNs offer a computationally efficient approach for inferring causality among a set of variables and have been applied to infer cellular networks^20^, protein-protein interactions^21^, protein-signalling networks^22^, gene regulatory networks^23,24^, and long-range intrachromosomal interactions^25^. However, inferring feasible BNs from data is challenging because the structure learning of BNs is sensitive to parameters and requires extensive prior knowledge to obtain expected structures.

Here, we present CIBER, a novel causal inference–based framework, for reconstructing cellular differentiation networks and identifying fate-determining features. CIBER provides a hybrid BN structure learning algorithm to detect information flow among cell states to learn a directed acyclic graph (DAG) representing the differentiation network. In addition to single-cell omics data which is critical for existing trajectory inference methods, CIBER can be applied to microarray or bulk sequencing data in which the cell states are discrete. We adopt the structural causal model^26,27^ to the learned DAG, utilize *in silico* perturbation as interventions for causal inference, and generate an effect matrix that quantifies the impacts of different features on each branch of the cellular differentiation network. Furthermore, we introduce differentiation driver feature (DDF) analysis to identify important features in the cellular differentiation process. DDF analysis can identify features crucial to the entire differentiation process or a specific lineage while not differentially expressed among lineages and can complement the regular differential feature analysis to help us understand the biological processes comprehensively.

We applied CIBER framework to study haematopoiesis using various types of omics or microarray data and obtained expected cell differentiation networks. The application on a mouse microarray dataset showed that CIBER can identify both expected and potential branches that disagreed with the conventional hierarchy while reported in previous studies. In this study, we use regular differential feature (hereafter named RDF) analysis to denote the analysis that identifies differentially expressed features directly from the cell-by-feature matrices representing gene expression or chromatin accessibility. In contrast to RDF, DDF analysis is performed by identifying important features based on the edge-by-feature effect matrices inferred via CIBER from the cell-by-feature matrices. DDF analysis enriches the techniques for identifying important features related to cellular differentiation process and can complement RDF analysis to help us understand biological processes more comprehensively. Through DDF analysis, CIBER identifies fate-determining features that are not deemed statistically significant in RDF analysis, can predict novel regulation associations and provide insight into the potential mechanism underlying the influence of TFs on biological processes. We conducted experiments on *Bcl11b*, a non-differentially expressed TF identified by CIBER, and validated CIBER’s prediction that it has a lymphoid-biased effect on the differentiation of haematopoietic stem cells.

## Results

### CIBER presents a novel framework for learning robust Bayesian network structures with minimal prior knowledge

Bayesian networks are directed acyclic graphical models that represent probabilistic dependencies between variables via edges and nodes^28^. The existing Bayesian network structure learning algorithms can be divided into three categories: constraint-based algorithms, score-based algorithms and hybrid algorithms^29,30^ (see Methods for details). One major issue in Bayesian network structure learning is that the obtained graph is highly sensitive to parameters and the observed data, and small changes in input may result in considerably different outputs. To obtain a well-constructed network, extensive prior knowledge is usually required to adjust the learned structures.

To overcome these limitations, we developed CIBER framework to generate robust Bayesian network structures given minimal prior knowledge, such that the proposed method can be generalized to investigate understudied biological processes. CIBER presents a new hybrid algorithm that uses a mixed strategy of resampling over single cells and combining the Bayesian networks learned within each resampling (Fig. 1). Stratified resampling is used to select random subsets of cells to construct the network, resulting in a set of networks with various structures. Specifically, for each subset of cells, centroids of different cell types are utilized to learn a Bayesian network, with each centroid representing the mean expression level of its corresponding cell type. The centroids are connected to generate an undirected skeleton according to Spearman’s correlation or the mutual information between them, with each edge representing the correlation between the two cell types it connects. Various algorithms for correlation estimation can be chosen, such as CMI2NI^31^ and SPACE^32^, under the support of GeNeCK^33^, an R framework for constructing gene regulatory networks. Based on the undirected skeleton, we utilized the hill climbing algorithm through the bnlearn^34^ package to search for the directed acyclic graph (DAG) with the highest Bayesian information criterion (Methods).

**Fig. 1.**
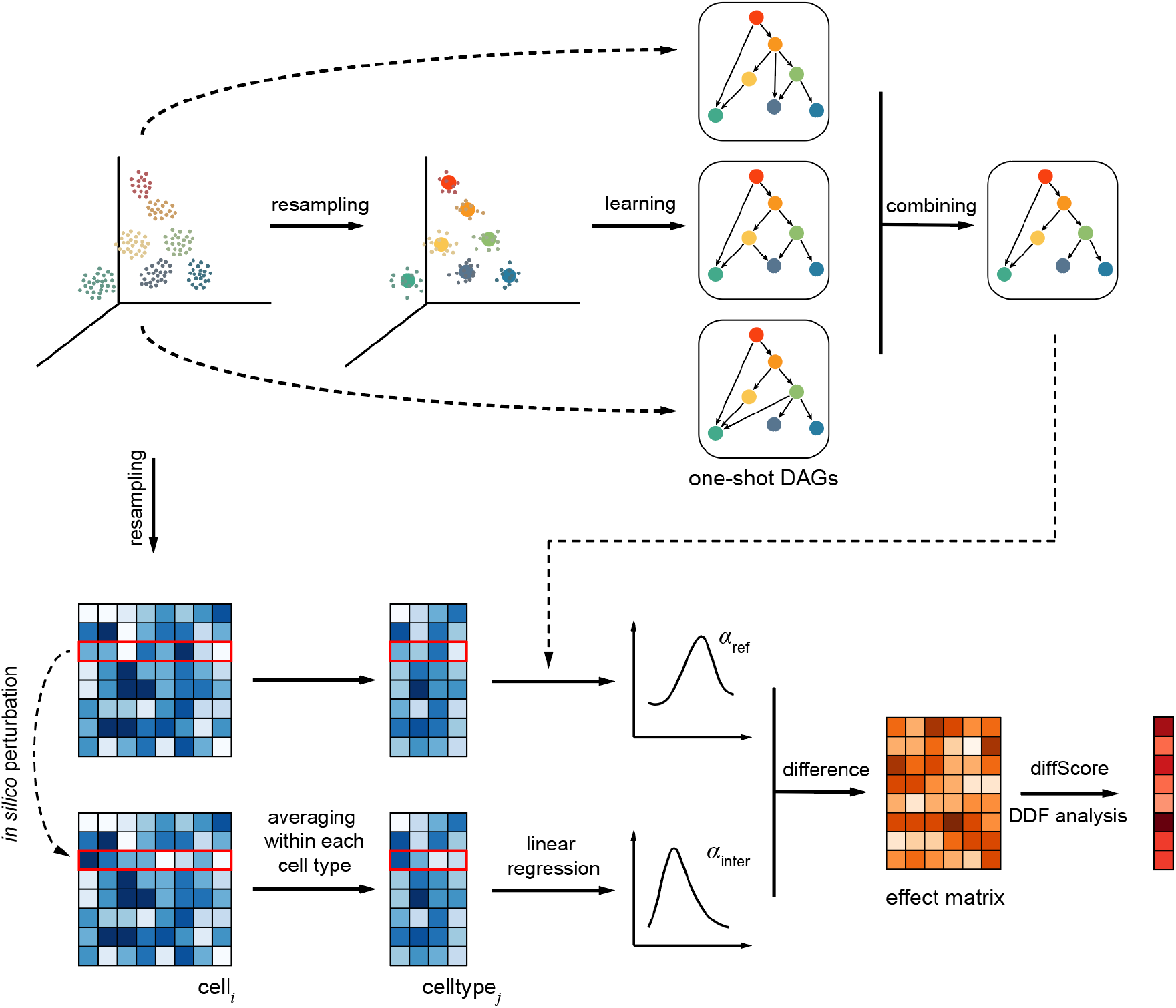
Simplified schematic of CIBER framework. The upper part shows the pipeline of the hybrid Bayesian network structure learning algorithm starting from cell resampling. Random subsets of cells are selected, and various DAGs are learned to represent the causation between the centroids of different cell types. Each DAG obtained within one resampling is referred to as a one-shot DAG for convenience in this study. Then, a combination strategy is applied over the one-shot DAGs to obtain the final Bayesian network. The bottom part shows the pipeline to generate effect matrix and to identify differentiation driving-genes. From the perspective of the structure causal model, a series of linear regressions are performed on the learned Bayesian network, with each child node taken as a dependent variable and its parent nodes taken as independent variables. A mixed strategy of resampling and interventions of *in silico* perturbation on a specific feature is utilized to obtain a pair of distributions for each edge, representing the reference and the intervened linear regression coefficients. The effect matrix is defined to quantify the difference between the two distributions and each element in the matrix reflects the effect of a feature on a differentiation branch. DDF analysis is performed on the effect matrix to identify fate-determining features.

The learned DAG is sensitive to the threshold used for evaluating correlations during the construction the undirected skeleton. In this study, we employed a range of sequential correlation thresholds, such as from 0 to 1 in steps of 0.01, to obtain a set of DAGs varying from dense to sparse for each resampling of cells. Approximate upper and lower boundaries for the number of edges in the network can be assigned according to the expected complexity of the network. DAGs with the number of edges within the specified boundaries are selected and combined to generate a directed graph, where each edge is weighted according to its frequency of appearance. The weight of each edge reflects its robustness, which in turn signifies the confidence level of the causation between the nodes it connects. However, the graph may not necessarily be acyclic, necessitating trimming and binarization to produce an unweighted DAG that retains the most robust structure. We refer to this DAG as the “one-shot DAG” since it is obtained within a single resampling. This process is then repeated several times to get a set of one-shot DAGs, over which a second combination is applied to generate a final DAG that captures the most consistent causation between variables (see Methods for detailed description). Our approach is applicable not only to single-cell omics data but also to microarray and bulk sequencing data, omitting the resampling step due to limited replicates.

### CIBER captures known differentiation structures in various datasets

We tested whether CIBER could obtain well-established cell lineage structures by applying it to study haematopoiesis using various types of data, including mouse microarray^35–37^, mouse bone marrow scRNA-seq^38^, human bone marrow scRNA-seq^39^ and human bone marrow scATAC-seq datasets^40^. The mouse microarray dataset contains comprehensive blood cell types, from which we selected a subset of representative types of progenitors to reconstruct the early haematopoietic hierarchy. The subset includes 11 cell types — haematopoietic stem cell (HSC), multipotent progenitor subset A (MPPa), multipotent progenitor subset B (MPPb), common myeloid progenitor (CMP), common lymphoid progenitor (CLP), granulocyte-macrophage progenitor (GMP), megakaryocyte-erythroid progenitor (MEP), megakaryocyte progenitor (MkP), precolony-forming unit erythroid (pCFU-E), B-cell-biased lymphoid progenitor (BLP) and double negative 1 thymocytes (DN1) — each containing 3 or 4 biological replicates. In this microarray dataset, the expression levels of 21,678 genes are detected by 45,101 probes^35–37^. If a gene is simultaneously detected by several probes, we set its expression level to the median among all the probes. Because of the stable expression levels over the limited size of replicates, we omitted resampling and performed the analysis on a set of fixed points that are the centroids obtained by averaging the expression level over biological replicates for each cell type.

The results revealed that CIBER can extract robust lineage structures and provide information on potential differentiation branches (Fig. 2b). Compared to the results obtained with a typical hybrid algorithm consisting of mutual information and hill climbing (hereafter named MI-hc), the CIBER-learned network was more consistent to the conventional haematopoietic hierarchy and more accurately captured the bifurcation structure (Fig. 2b, c). In these figures, the edges contradicting prior knowledge are coloured in red. The CIBER-learned structure is robust without the need to manually set the exact correlation threshold, unlike MI-hc, which is sensitive to parameters, and small changes in the correlation threshold can result in considerably different structures (Extended Data Fig. 1). In addition to the well-known branches of the HSC → MPPa/MPPb→lymphoid/myeloid lineages and those inside the lymphoid or myeloid lineage, CIBER identified three unconventional interlineage edges that were reported to be possible in previous studies, CLP→GMP^41^ and CMP→BLP/DN1^42,43^.

**Fig. 2.**
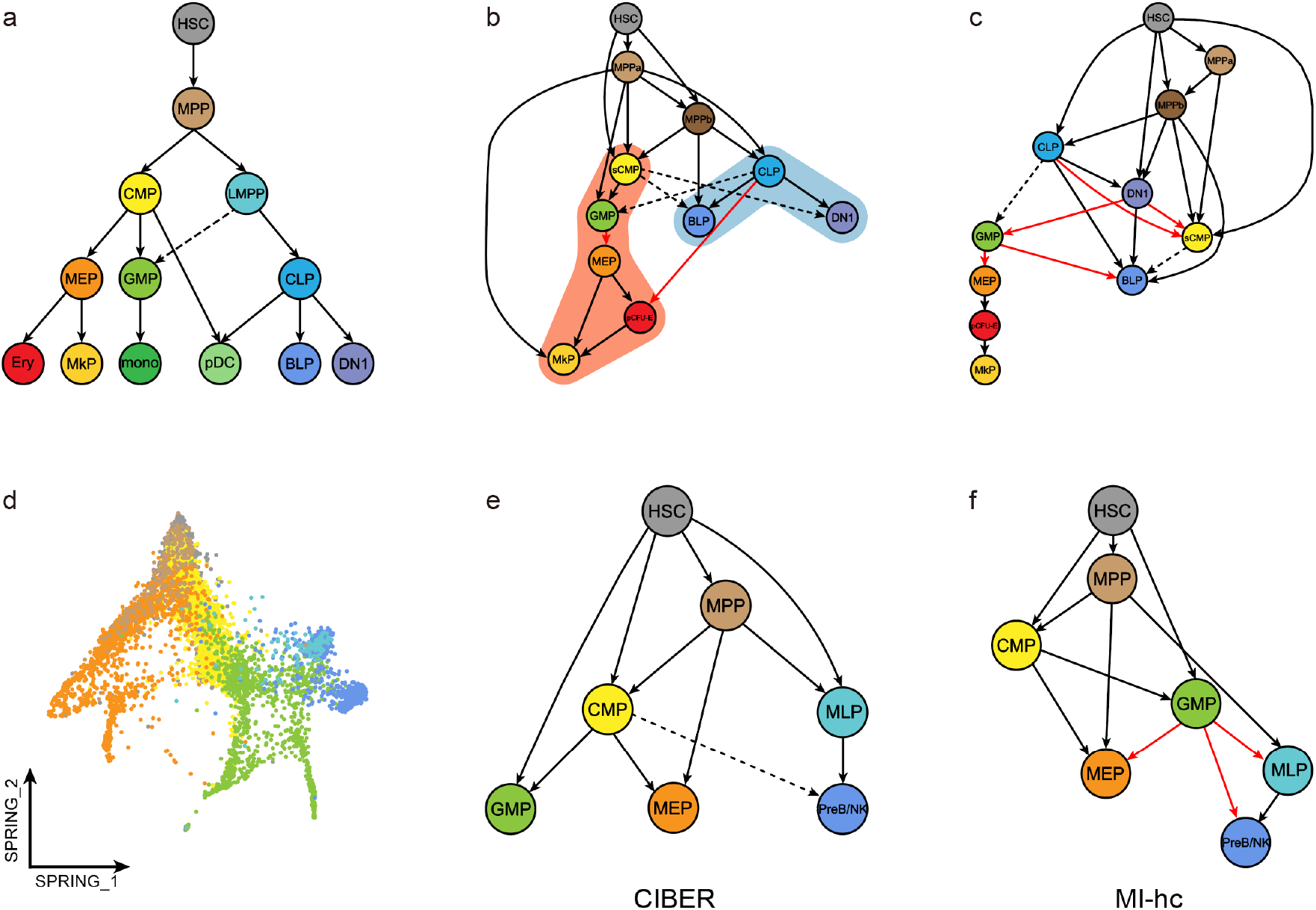
CIBER captures conventional differentiation structures in various datasets. **a**, Conventional haematopoietic hierarchy. **b**, CIBER result of the mouse microarray dataset, with the blue/red-shaded part representing the lymphoid/myeloid lineage. **c**, Best-tuned structure learned with MI-hc, a typical hybrid algorithm consisting of mutual information and hill climbing, using the same dataset as **b**; the structures learned under other correlation thresholds are shown in Extended Data Fig. 1. The dashed edges represent unconventional interlineage branches that were reported to be possible in previous studies, while the red edges represent less credible branches. **b** and **c** show that the CIBER result is consistent with conventional knowledge with a larger number and a higher percentage of edges and captures the bifurcation structure more accurately than MI-hc. **d-f**, Low-dimensional landscapes^39^, the CIBER and MI-hc results of the human scRNA-seq dataset. Results for more datasets are shown in Extended Data Fig. 2.

We also applied CIBER to single-cell omics datasets, following the entire pipeline, including the resampling procedure. The structures learned by CIBER again identified larger numbers and higher percentages of conventional edges compared to the results of the typical hybrid algorithm MI-hc (Fig. 2d-f, Extended Data Fig. 2). Furthermore, the cell types in CIBER-learned structures were in relative positions consistent with the low-dimensional landscapes of diffusion maps^38^, SPRING^39^ or PCA^40^ obtained in previous studies.

### CIBER quantifies the effects of features on the cellular differentiation hierarchy

CIBER enables quantification of the effects of features on the cellular differentiation hierarchy by utilizing the structural causal model (SCM), a conventional causal inference framework that combines probabilistic graphical models with the notion of interventions^26,27^. Similar to Bayesian networks, SCM considers a set of associated observables in the form of a directed acyclic graph while viewing each node as a function of its parents in the graph. A natural restriction of SCM is the assumption of a Gaussian linear additive noise model, which limits the functions connecting each node to its parents to the form of linear regression (see Methods). In the SCM framework, interventions can be performed by changing the form of particular functions or the distribution of specific nodes to infer the effect of causality.

Here, we introduced the concept of SCM into the directed acyclic graph learned by CIBER and performed causal inference by applying *in silico* perturbation to the expression of specific features as interventions (Fig. 1). Specifically, we performed feature permutation on single-cell omics data and feature deletion on microarray or bulk sequencing data. We then recorded the coefficient associated with each edge in the Gaussian linear additive SCM, with and without interventions performed on specific features, and defined an effect matrix to measure the change in the coefficient due to the interventions. The effect matrix quantifies the impacts of the features on each edge in the cellular differentiation network, with positive values implying a possible promoting function on the corresponding differentiation branch and negative values implying a possible inhibiting function.

### Differentiation driver feature (DDF) analysis identifies features determining cell fate

We present a novel method, called DDF analysis, which distinguishes itself from regular differential feature (RDF) analysis by identifying important features that are crucial to the cellular differentiation process via effect matrices obtained by *in silico* perturbation. The method quantifies the impact of individual features on the cellular differentiation network or specific lineages through diffScores (see Methods). The diffScore of a feature on the entire network is defined by summarizing the absolute values related to this feature in the effect matrix, while the diffScore on a specific lineage is defined as the sum of the absolute values related to the edges pointing to cell types in the lineage subtracting the original values of the edges pointing from the lineage to other lineages. Our results show that DDF analysis can identify crucial features which are not differentially expressed between lineages, and provides a paradigm to systematically investigate a feature’s impact on the cellular differentiation process even in the absence of some important cell types. Additionally, DDF analysis can be combined with RDF analysis to provide a comprehensive understanding of biological processes due to the heterogeneous information captured by the two methods.

To evaluate the effectiveness of DDF analysis, we applied it to the mouse bone marrow scRNA-seq dataset and compared the results with that of the RDF analysis obtained by ranking genes according to the p value of Wilcoxon’s signed-rank tests. To compare the results of the two analyses, we selected the same number of top genes on the two lists, and conducted a literature search on the top-ranked genes (Fig. 3a-c). Results showed that the RDF and DDF analyses both captured a part of the genes that have been proven to be important in haematopoiesis: a few important fate-determining genes identified by RDF analysis, such as *Gata2*^44^, were not captured by DDF analysis; on the other hand, DDF analysis identified important genes that were not differentially expressed between the lymphoid and myeloid cell types, such as *Mpl*^45^, *Ccl9*^46^ and *Elane*^47^; nevertheless, some genes, such as *Flt3*^48^, *Cd34*^49^ and *Dntt*^50^, were identified by both analyses.

**Fig. 3.**
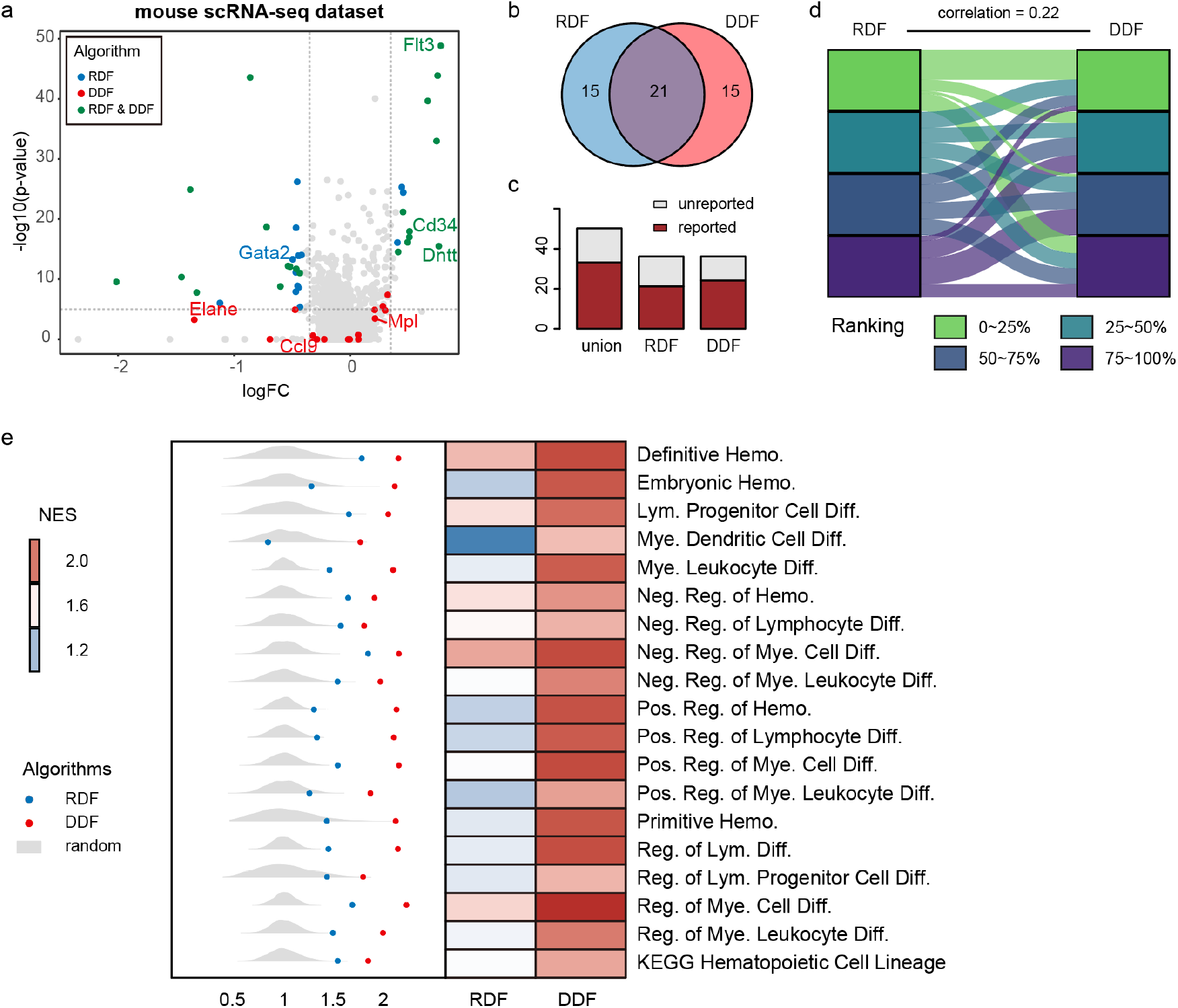
DDF identifies fate-determining features. The plots show the results on RDF and DDF gene lists obtained with the mouse scRNA-seq dataset. **a**, Features are allocated by log fold change and log p value obtained in the RDF analysis, from which the top differentially expressed genes are coloured, as well as the same number of DDF-selected features important to the entire network. **b**, The number of shared top features between the two feature sets. **c**, Literature search result shows that the DDF and RDF analyses identify different important features, in which the red parts represent the reported features. **d**, Correlation between the ranked lists shows that the DDF analysis captures different information from the RDF analysis. The tiles represent the top 25%, 25∼50%, 50∼75% and bottom 25% features in RDF or DDF ranking lists. **e**, GSEA normalized enrichment scores (NES) with terms related to haematopoiesis or haematopoietic differentiation in the GO and KEGG databases. DDF list obtains higher NES than RDF on all the terms. Dots in the left part represent the NES of RDF and DDF ranking lists, while the distributions show the background NES given randomly ranked gene lists. (Hemo., Hemopoiesis; Diff., Differentiation; Lym., Lymphoid; Mye., Myeloid; Reg., Regulation; Pos., Positive; Neg., Negative.)

We calculated Spearman’s correlation between the two gene lists and found that they were weak correlated (Fig. 3d), and further evaluated their behaviours on enriching important genes related to haematopoiesis or haematopoietic differentiation. Specifically, we calculated GSEA normalized enrichment scores (NES) for gene lists with respect to all the GO and KEGG terms related to haemopoiesis or haematopoietic differentiation (Fig. 3e). GSEA NES results showed that DDF surpassed RDF on almost all these pathways. Note that the dataset did not contain any lymphoid-lineage cell types other than lymphomyeloid-primed progenitor (LMPP), an early lymphoid-biased progenitor. This illustrates that DDF analysis performs well even in the absence of important cell types, due potentially to the fact that it examines the entire cellular differentiation network rather than being restricted to the difference between subgroups of cells, as RDF analysis did.

Next, we applied the same analyses to the human scRNA-seq dataset and mouse microarray dataset. Similar results were obtained for the human scRNA-seq dataset to the results of mouse scRNA-seq dataset described above (Extended Data Fig. 3). The microarray dataset offered us an opportunity to study the behaviour of DDF analysis in scenarios where most cell types are available and are therefore friendly to RDF analysis. The result showed the RDF list performed slightly better than the DDF list in 12 out of 22 terms. However, the literature search still showed that DDF analysis identified important genes that are not differentially expressed, demonstrating that DDF analysis can complement RDF analysis to better identify genes crucial to cell differentiation (Extended Data Fig. 4).

### CIBER identifies crucial TFs regulating biological processes

We then applied CIBER and DDF analysis to a human bone marrow scATAC-seq dataset^40^, and identified TFs crucial to haematopoiesis based on deviation score matrix calculated with chromVAR^51^. RDF analysis was also performed to identify the TFs exhibiting significant differences in deviation score between lymphoid and myeloid cell types. The top TFs identified by DDF alone include *SPI1, SPIB* and *CEBPB*; whereas TFs like *TCF3, LYL1* and *EBF1* were exclusively identified by RDF analysis. Both analyses uncovered TFs such as *GATA1, GATA2, GATA3, ID3* and *LMO2* (Fig. 4c).

**Fig. 4.**
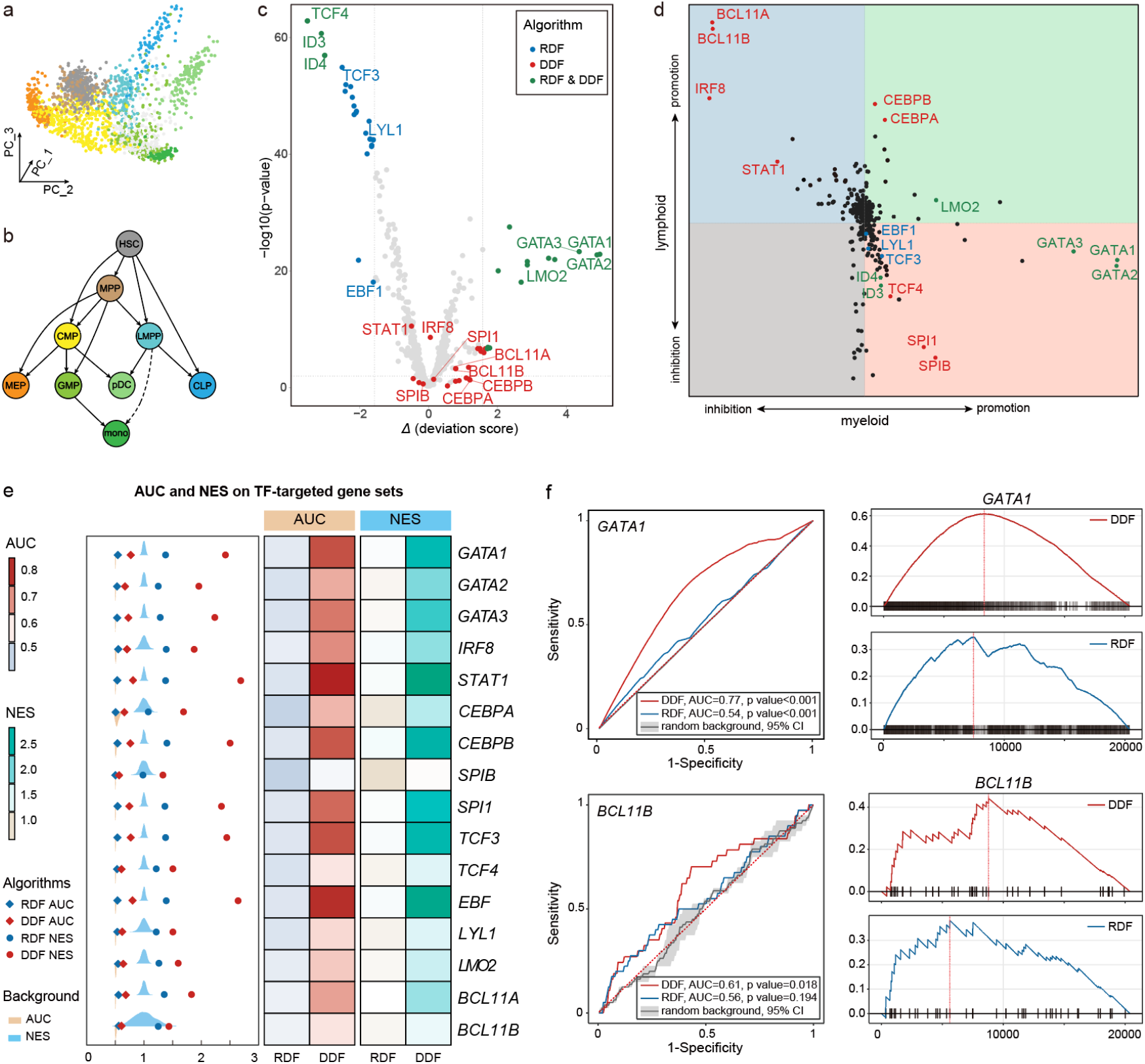
CIBER identifies driver TFs of biological processes. **a**, PCA reduction of a human bone marrow scATAC-seq dataset^40^. **b**, Differentiation hierarchy learned with CIBER. **c**, Top TFs identified by RDF and DDF analyses. Among them, several reported important TFs^40^ are annotated. **d**, Lineage-specific diffScores quantify the effects of each TF on haematopoiesis under a higher resolution of promoting or suppressing specific lineages. **e**, Comparison of rankings obtained with DDF and RDF analyses using ROC or GSEA analyses applied with the set of genes regulated by each TF. AUC and GSEA NES results show DDF surpasses RDF on each gene set and significantly surpasses random background; background is set as the distribution of 1,000 random gene rankings. **f**, AUC and NES Results on *GATA1*-targeted and *BCL11B-*targeted gene sets respectively. For *GATA1-*targeted gene set, both DDF and RDF significantly surpass random background, and DDF scores higher; for *BCL11B-*targeted gene set, DDF significantly surpasses random background, while no statistical difference exists between RDF and the background (DDF NES p value=0.009, RDF NES p value=0.103, DDF AUC p value=0.018, RDF p value=0.194). Thus, compared with regular differential analysis, CIBER can better identify important TFs and enrich reported regulated genes, providing insight into potential regulated genes through which a TF influences the process of haematopoiesis.

With lineage-specific diffScores, DDF analysis can further predict the potential effects of each TF on promoting or inhibiting specific lineages (Fig. 4d). In addition to the previously reported TFs, we noted that DDF analysis exclusively identified *BCL11B*, and predicted that it may have similar impacts on haemopoiesis of promoting the lymphoid lineage while inhibiting the myeloid lineage differentiation with previous reported *IRF8*^52^ and *BCL11A*^53,54^.

We further examined if the target genes of the TFs identified in the scATAC-seq dataset were also enriched in the DDF or RDF analysis of an independent human bone marrow scRNA-seq dataset since TFs regulate cellular function by orchestrating the transcription of downstream target genes. To achieve this, we collected target genes for 15 reported TFs^40^ and *BCL11B* from 5 databases (TRRUST^55^, JASPAR^56^, ChEA^57^, Motifmap^58^ and ENCODE^59^) and ranked the target genes via DDF or RDF analysis performed on the scRNA-seq dataset. To measure the enrichment of TF-targeted genes, we calculated two types of metrics, the normalized gene set enrichment score (NES) and the area under the receiver operating characteristic curve (AUC) (Fig. 4e, Methods). When calculating AUC, the receiver operating characteristic (ROC) curve was drawn by assigning TF-targeted genes as positive and all other genes as negative samples. Background distributions of NES and AUC were obtained with 1,000 randomly shuffled ranking lists of all the genes and were used for calculating p value. Results show that under both metrics, DDF list surpasses RDF list on all gene sets targeted by different TFs (Fig. 4e, Extended Data Fig. 5). Therefore, CIBER can predict novel regulation associations and provide insight into the potential mechanism underlying the influence of TFs on biological processes. For example, for the *GATA1-*targeted gene set, DDF list scores higher than RDF list, and both lists significantly surpass the background; in the case of *BCL11B*, only the DDF list significantly surpasses the background (DDF NES p value=0.009, RDF NES p value=0.103, DDF AUC p value=0.018, RDF p value=0.194) (Fig. 4f).

### In vivo experiments validate *BCL11B* as a crucial TF to fate determination in haematopoiesis

The above mentioned results indicate that the TF *Bcl11b* may influence the differentiation of haematopoietic stem cells. To validate this, we constructed *Bcl11b*-deficient mice (*Bcl11b*^flox/flox^, Vav-iCre^+^), hereafter named *Bcl11b*^-/-^ mice, by specifically knocking out *Bcl11b* in the blood cells. Results showed that the lifespan of *Bcl11b*^-/-^ mice was significantly shorter than that of wild-type (WT) mice in the same cohort, indicating that *Bcl11b* is essential for the maintenance of hematopoietic system (Fig. 5a). To examine the haematopoietic dynamicity of *Bcl11b*^*-/-*^ mice, we treated *Bcl11b*^*-/-*^ and WT mice with 5-fluorouracil (5-FU) and analyzed the lineage composition in peripheral blood (Fig. 5b). The result showed that myeloid cells were significantly reduced in both groups five days after 5-FU injection, indicating that they are sensitive to DNA damage challenge (Fig. 5c). Notably, T cells were almost undetectable in *Bcl11b*^*-/-*^ mice, while remaining static in the WT group five days after 5-FU injection compared to non-treated mice, suggesting that the *Bcl11b* plays an essential role in maintaining T cell in response to DNA damage. Between days 10 to 20, we observed a significant decrease in the percentage of both T and B lineages compared to WT mice, indicating that *Bcl11b* is crucial for lymphoid differentiation, consistent with our prediction (Fig. 4b).

**Fig. 5.**
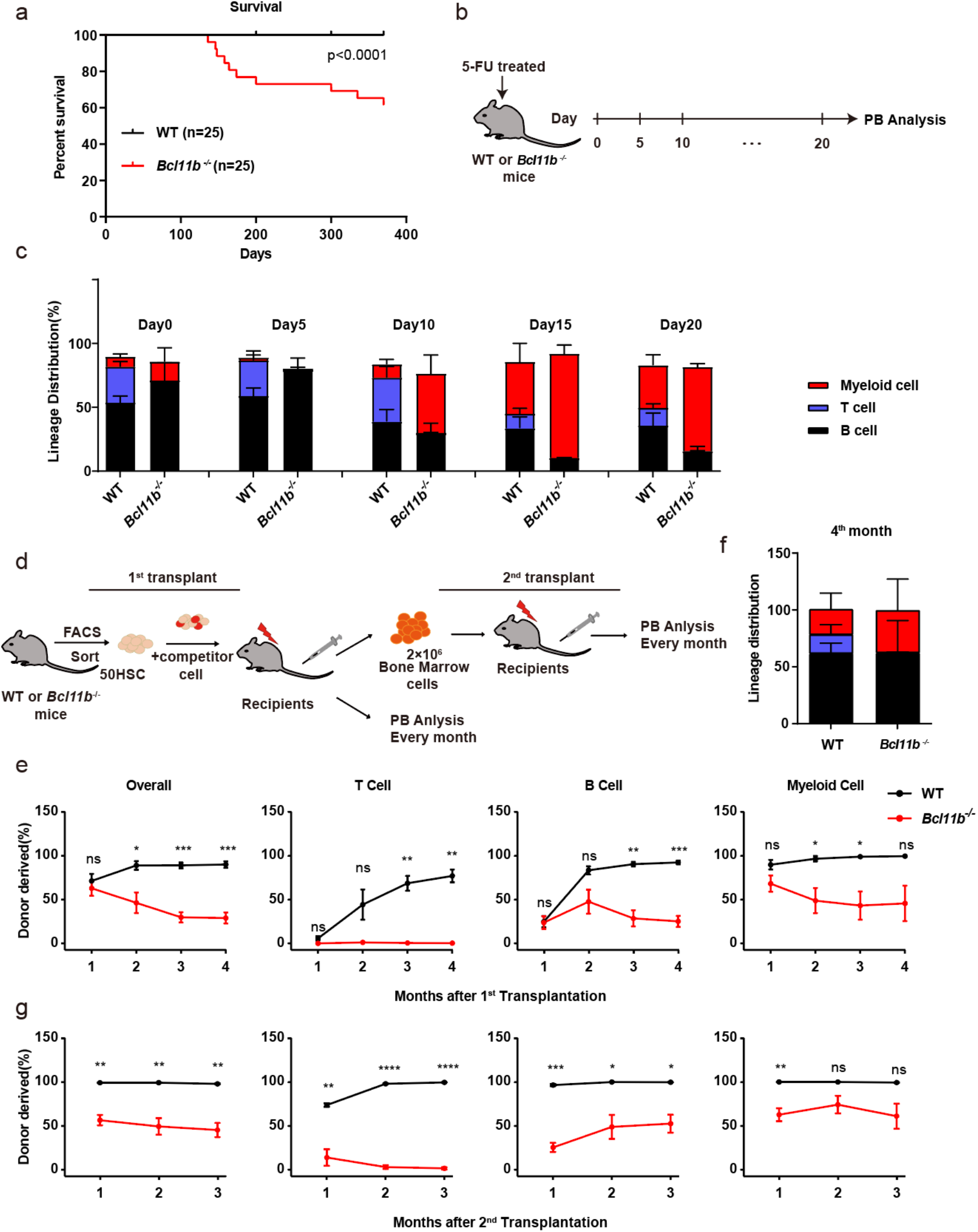
Experimental validation of CIBER predictions for *Bcl11b*. **a**, Results of survival experiments of WT and *Bcl11b*^-/-^ mice in their native state show the survival capacity of *Bcl11b*^-/-^ mice is hampered. Experiments (**b**) and Lineage distribution (**c**) of reconstituted B cells, T cells and myeloid cells in peripheral blood of WT and *Bcl11b*^-/-^ mice after 5-FU treatment. **d**, Scheme of the transplantation strategy for *Bcl11b*^-/-^ mice. Fifty HSCs freshly isolated from 2-month-old WT or *Bcl11b*^-/-^ mice were transplanted into lethally irradiated recipients along with 3×10^5^ competitor cells. For the 2^nd^ transplants, 2×10^6^ whole bone marrow cells were transferred from primary recipient mice. Chimera in peripheral blood was examined every month for three months. **e**, Contributions of donor cells to peripheral blood overall (CD45.2^+^), B cells (B220^+^), T cells (CD3^+^) and myeloid cells (Mac-1^+^) every month after 1^st^ transplantation. (n=4-5 mice per group. Data are shown as the mean ± SD.) **f**, Lineage distribution of B cells, T cells, and myeloid cells among donor-derived cells in peripheral blood four months after transplantation. (n=4-5 mice per group. Data are shown as the mean ± SD.). **g**, Contributions of donor cells to peripheral blood overall (CD45.2^+^), B cells (B220^+^), T cells (CD3^+^) and myeloid cells (Mac-1^+^) every month after 2^nd^ transplantation. (n=4-5 mice per group. Data are shown as the mean ± SD.)

To assess the impact of *Bcl11b*-deficiency on the long-term haematopoietic reconstitution capacity and differentiation of HSCs, we conducted competitive transplantation assays by injecting 50 freshly isolated HSCs either from *Bcl11b*^-/-^ or WT mice into lethally irradiated recipients together with 3×10^5^ total bone marrow cells as competitor (Fig. 5d). Peripheral blood chimerism was monitored monthly until the 4th month. Our results demonstrate that the targeted dysfunction of *Bcl11b* severely impairs the reconstitution capacity of HSCs (Fig. 5e). Notably, the differentiation potential of *Bcl11b*^*-/-*^ HSCs to T lineage is severely compromised from the 2nd month after transplantation. By the end of the 4th month, *Bcl11b*^*-/-*^ HSCs exhibit differentiation bias towards the myeloid lineage at the expense of T lineage (Fig. 5f), which is consistent with our prediction (Fig. 4b).

To further investigate the influence of *Bcl11b* dysfunction on the self-renewal capacity of HSCs, we performed secondary transplantation assays (Fig. 5d). Our findings demonstrate that the chimera of *Bcl11b*^*-/-*^-derived cells in the peripheral blood of recipients is significantly lower than that of the control group, including B, T and myeloid lineages (Fig. 5g), suggesting that the self-renewal capacity of *Bcl11b*^*-/-*^ HSCs is impaired. In summary, our results indicate that *Bcl11b* plays a critical role in promoting T lymphocyte differentiation and that its dysfunction impairs the reconstitution and self-renewal capacity of HSCs, resulting in a differentiation bias towards the myeloid lineage.

## Discussion

Recent advancements in single-cell omics data have allowed researchers to study developmental pathways and lineage relationships with unprecedented resolution. However, most trajectory inference methods are based on expression pattern similarities and provide cell fate probabilities, which may not accurately reflect the complex causation-driven phenomena of cellular differentiation. In this study, we present CIBER framework based on causal inference to reconstruct cellular differentiation networks and identify fate-determining features, which provides a novel paradigm for dissecting cell state transitions other than trajectory inference and differential analysis.

Compared to trajectory inference methods, CIBER has several advantages. It is applicable under a broader range of conditions, including datasets with limited cells or cell types, and even those obtained by microarray or bulk sequencing techniques. Furthermore, unlike most similarity-based methods, CIBER provides a causation-based perspective on the cellular differentiation process.

CIBER providess a robust Bayesian network structure learning strategy to obtain feasible structures with minimal prior knowledge. The applications of CIBER show that it is suitable to different types of datasets and that it may help in identifying potential differentiation branches. Based on the concept of the structural causal model and interventions of *in silico* feature perturbation, CIBER builds an effect matrix quantifying the impacts of features on differentiation branches and can be used for identifying fate-determining features through DDF analysis. DDF analysis can identify features crucial to the entire differentiation process or a specific lineage while not differentially expressed among lineages and can complement the regular differential feature analysis to help us understand the biological processes comprehensively. Besides, DDF analysis is effective in scenarios even in the absence of important cell types.

Despite its advantages, CIBER has some limitations. CIBER may be time-consuming, and should take the centroid of cell types or clusters as nodes rather than cells, and it may not be suitable for extremely complex networks with too many nodes and edges. We found that CIBER was most effective for hierarchical structures with approximately 5 layers and less than 20 nodes.

In conclusion, the CIBER framework provides a causal perspective for the reconstruction of cellular differentiation networks and the evaluation of how features influence differentiation branches. DDF analysis enriches existing techniques for identifying important features and can be combined with differential analysis to help us understand biological processes more comprehensively. Overall, CIBER has the potential to revolutionize the study of cellular differentiation and provides insights into the underlying mechanisms of developmental pathways.

## Methods

### Data preprocessing

Normalized mouse microarray, mouse bone marrow scRNA-seq, human bone marrow scRNA-seq and human bone marrow scATAC-seq datasets were obtained from the data sources. In the microarray dataset, each cell type had 3 or 4 biological replicates, and the expression levels of 21,678 genes were detected by 45,101 probes. For each gene detected by multiple probes, the median value was adopted as its expression level. The top variable features were first selected for Bayesian network structure learning; 3,000 features were selected in the mouse microarray dataset, 5,000 in the mouse bone marrow scRNA-seq dataset, 4,000 in the human bone marrow scRNA-seq dataset and 1,764 (all TFs) in the human bone marrow scATAC-seq dataset.

### Construction of cellular differentiation networks

The existing Bayesian network structure learning algorithms can be divided into three categories: constraint-based algorithms, score-based algorithms and hybrid algorithms^29,30^. Constraint-based algorithms construct a Bayesian network by first generating an undirected skeleton via Markov blanket^60^ or correlation and then orienting the skeleton under acyclicity constraints to obtain the directed acyclic graph (DAG) that best satisfies the conditional independence test. The effectiveness and efficiency of constraint-based algorithms are limited due to the lack of robustness and the high complexity of the conditional independence test. Score-based algorithms evaluate how well a DAG fits the observed data with a score such as the Akaike information criterion^61^ or Bayesian information criterion^62^, with the goal of identifying the graph obtaining the highest score. Major issues for score-based algorithms include the exponentially growing search space as the number of variables increases and the search algorithm may lead to a local optimum. One solution to the limitations of the abovementioned two types of algorithms is hybrid algorithms, which first generate an undirected skeleton and then search for the DAG with the highest score among the structures restricted to this skeleton. However, the three types of BN structure learning face a common problem: they are sensitive to observations and parameters, and small changes in input can lead to considerably different outputs. Hence, considerable prior knowledge is needed to tune and obtain expected structures.

To overcome these limitations, CIBER employs several strategies to ensure a well-structured hierarchy with minimal prior knowledge of assigning credible upper and lower boundaries for the number of edges in the network and setting stem cells as the root. CIBER utilizes a multistep strategy to obtain robust BN structures — resampling, undirected skeleton learning, DAG learning, combining DAGs and trimming branches. Particularly, for datasets with limited biological replicates, such as microarray datasets, the resampling step is omitted. By combining the DAGs obtained with different resampled subsets of single cells and various parameters, we can extract the most robust associations between variables, which have a higher probability of being the true causation. The details of the steps are described below.

### Resampling strategy

The resampling strategy maximizes the efficacy of large single-cell omics datasets. A stratified resampling strategy was performed for the single-cell omics datasets to generate a subset of cells, while this strategy was omitted for the microarray dataset. For the microarray dataset and each subset of single cells after resampling, we calculated the mean expression level of each cell type to obtain a mean expression matrix, with each column representing a cell type and each row representing a specific feature. The resampling ratio can be set to approximately 0.3 to prevent high randomness or homogeneity between the obtained subsets. Similarly, the cell number of each cell type was suggested to be greater than 50 to prevent random effects.

### Undirected skeleton learning

With the mean expression matrix as an input, an undirected skeleton was built to describe the correlation between each pair of cell types, with each node representing a cell type and each edge representing the correlation between the connected cell types. Various metrics can be utilized to evaluate the correlation, such as Spearman’s correlation or the mutual information. In addition to the correlation, several algorithms developed for undirected skeleton construction can be conveniently chosen under the support of GeNeCK^33^. The default algorithm used in CIBER is CMI2NI^31^, an approach that was developed for reverse-engineering gene regulatory networks by quantifying the conditional mutual inclusive information between two genes given a third gene by calculating the Kullback–Leibler divergence. The undirected skeleton is determined by the threshold used for judging the correlation, and a sparser skeleton will be obtained given a higher correlation threshold. A set of sequential correlation thresholds was used to obtain a series of undirected skeletons for further DAG learning.

### DAG learning

Typically, two causally associated variables should also be correlated. The undirected skeleton provides us with a reference that can be used to determine the causal DAG. We excluded absent edges in the undirected skeleton in the alternative DAGs and utilized the existing score-based structure learning algorithm offered in the bnlearn^34^ package to obtain the DAGs. The hill climbing algorithm was used to search for the DAG with the highest Bayesian information criterion (BIC)^62^. BIC is commonly used in Bayesian network structure learning to evaluate how well a graph *G* fits the observed data *D* and can be formulated as follows:

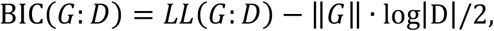

where *LL*(*G*: *D*) describes the likelihood that *G* fits *D*, and ‖*G*‖ and log|D| represent the complexity of the graph structure and dataset, respectively. Thus, hill climbing seeks to identify the least complex DAG that best explains the observed data.

### Combination of DAGs and trimming branches

A DAG can be generated with each correlation threshold under a resampled subset of cells. By changing the correlation threshold, we can obtain a series of different DAGs. The DAGs with the number of edges located between the prior-set upper and lower boundaries are selected and combined to obtain a directed graph, with each edge weighted according to its frequency of occurrence. According to the assumption that the true causation should be robust to different parameters, a combined weighted DAG can be generated by extracting the highest weighted edges and removing the lowest weighted edges in cycles.

Then, this DAG is binarized to obtain an unweighted DAG named one-shot DAG in this study, which indicates the robust causation within this batch of cells. For microarray or bulk sequencing datasets, the one-shot DAG is the final learned Bayesian network structure. However, for single-cell omics datasets, the one-shot DAG reflects only the possible causation within a specific subset of the whole dataset. Therefore, a similar combination is applied over different one-shot DAGs to generate a quasi-final DAG. Under some special conditions, when generating the quasi-final DAG, additive prior knowledge may be needed to determine which edge should be severed if two edges with the same weights occur in one cycle. The quasi-final DAG may contain some edges that are not robust while being kept for the reason that their opposite edges occur even less.

An optional procedure can be performed on the quasi-final DAG to limit the maximum number of parents for each node and to remove redundant low-weighted edges. Specifically, under the structural causal model framework, we fit the quasi-final DAG to the mean expression matrix of all the cells in the original dataset twice, once with the top 25% variable features and once with the top 25∼50% variable features, and maintain the edges that remain stable while removing edges with low-weighted coefficients.

### Construction of the effect matrix

Based on the learned Bayesian network, the SCM framework can be utilized to build an effect matrix that quantifies the effects of different features on various differentiation branches. SCM is a conventional causal inference framework that combines probabilistic graphical models and the notion of interventions. This framework considers a set of associated observables with the nodes of a directed acyclic graph while viewing each node *X*_*i*_ as a function of its parents **PA**_*i*_ in the graph:

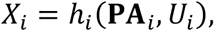

where *U*_*i*_ represents noise or exogenous variables. Under a linear additive noise assumption, *X*_*i*_ can be viewed partially as a linear regression of **PA**_*i*_:

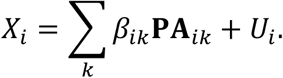

Interventions are performed to infer causality based on the SCM equations. Typically, by intervening in the form of a function or distribution of **PA**_*i*_, a latent expression of *X*_*i*_ can be obtained. The difference between the observed and latent expressions of *X*_*i*_ is measured to estimate the effect of causality.

In the construction of the effect matrix, we perform interventions by *in silico* perturbation of permuting (or deleting) the expression of a specific feature in all cell types and estimate the effect of causality by concentrating on the change in *β*_*ik*_ instead of the change in *X*_*i*_. The coefficient *β*_*ik*_ represents the association between two connected cell types and reflects the information flowing from **PA**_*ik*_ to *X*_*i*_ and the information of the corresponding differentiation branch. Thus, the change in the coefficients caused by intervening in a specific feature quantifies the effect of that feature on the cellular differentiation process.

For the single-cell omics dataset, we used a resampling strategy similar to that performed in reconstructing networks to record the distribution of the coefficient associated with a particular edge *e*_0_ obtained without and with interventions on a specific feature *f*_0_, which we termed *α*_ref_ and *α*_inter_, respectively, where *α* represents the abovementioned *β*_*ik*_. For one resampling and intervention, the matrix EffectMat_sampling_ is defined as

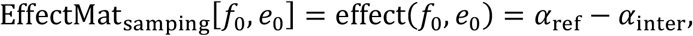

where EffectMat_sampling_ [*f*_0_, *e*_0_] is the row *f*_0_ and column *e*_0_ element in the matrix, and effect(*f*_0_, *e*_0_) is the effect of feature *f*_0_ on edge *e*_0_. By resampling and permuting several times, we obtain a series of EffectMat_sampling_ matrices, which can be horizontally merged to generate a final effect matrix in which the rows consist of all features and the columns consist of all edges multiplied by the resampling and permutation times. For microarray or bulk sequencing datasets, the effect matrix is constructed with the same procedures, except that the resampling step is omitted and the intervention is performed with deletion instead of permutation.

### DDF analysis

We developed DDF analysis to identify features important to specific lineages or the entire cellular differentiation network according to the effect matrix. When applying DDF analysis, diffScores are defined to evaluate the effects of different features on the differentiation process or a specific lineage.

For the haematopoiesis network, we defined diffSum, diffScore_lym_ and diffScore_myl_ to quantify the effects of the features on the entire haematopoietic hierarchy, the lymphoid-lineage and the myeloid-lineage, respectively. Let *e* represent an edge; the effect of feature *f*_0_ on the entire haematopoietic cellular differentiation network *G* is defined as

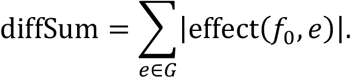

The lymphoid-lineage and myeloid-lineage effects are defined as

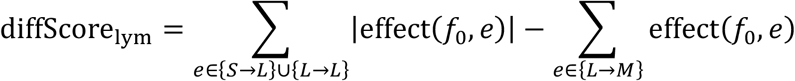

and

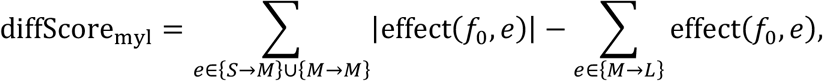

where *S* represents early cell types, including stem cells and multipotent cells, *L* represents lymphoid-lineage cell types, *M* represents myeloid-lineage cell types, and {*S* → *L*} represents the set of edges pointing from cell types in set *S* to cell types in set *L*.

### Transcription factor–targeted gene set GSEA and ROC analyses

For each TF, we collect the regulated genes from 5 databases (TRRUST, JASPAR, ChEA, Motifmap and ENCODE) and apply GSEA and receiver operating characteristic (ROC) analyses on this gene set. GSEA and ROC analyses are performed with the ranked gene lists obtained by DDF and RDF analyses.

Specifically, normalized enrichment score (NES) is calculated in GSEA analysis. When applying ROC analysis, we plot ROC curve by assigning the TF-targeted genes as positive and the others as negative samples. In ROC curve, the true positive rate is referred to as sensitivity and the true negative rate is referred to as specificity,

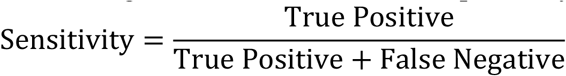

and

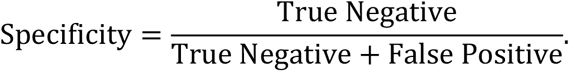

AUC score is calculated as the area under ROC curve.

Background distributions are obtained by calculating NES and AUC of 1,000 randomly ordered lists of all the genes.

### Competitive HSC transplantation and analysis

In all experiments (unless otherwise specified), mice were homozygous for the *Bcl11b* allele. *Bcl11b*^-/-^ (C57BL/6, CD45.2) mice, C57BL/6-SJL (CD45.1) (Stock No: 002014) and C57BL/6 WT (CD45.2) (Stock No: 000664) mice were obtained from the Jackson Laboratory. CD45.1/2 mice were heterozygotes from CD45.1 and CD45.2 mice. All mice were kept in specific-pathogen-free (SPF), AAALAC-accredited animal care facilities at the Laboratory Animal Research Center, Tsinghua University and all procedures were approved by the Institutional Animal Care and Use Committee of Tsinghua University. Bone marrow cells were harvested by crushing the femur and tibias with pestle and mortal in Hanks balanced salt solution (HBSS) with 2% fetal bovine serum and 1% N-2-hydroxyethylpiperazine-N’-2-ethanesulfonic acid buffer (HBSS+). Cells were stained with antibodies labelled with fluorochromes. For hematopoietic stem and progenitor cells enrichment, bone marrow cells were stained with c-Kit-APC. Hematopoietic populations were identified by flow cytometry using BD LSRFortessa (BD Biosciences). Data were analysed using FlowJo software.

For the primary competitive HSC transplantation assay, 50 freshly isolated HSCs from WT or *Bcl11b*^-/-^ mice (CD45.2) were transplanted into lethally irradiated (10 Gy) recipients (CD45.1/2) together with 3×10^5^ competitor cells (CD45.1). Donor-derived chimerism (including B cells, T cells and myeloid cells) in the peripheral blood of recipients was analysed at monthly intervals.

Two-tailed unpaired Student’s *t* test was applied for analysing experiment data after testing for normal distribution. All histograms were plotted by GraphPad Prism 8 software, and p value <0.05 was regarded as significant for all tests. All data are depicted as the mean ± SD.

## Data availability

The datasets used in this work were downloaded. Mouse microarray dataset^35–37^ can be found at GEO (GSE34723). Mouse bone marrow scRNA-seq data^38^ can be found at GEO (GSE81682). Human bone marrow scRNA-seq data^39^ can be found at GEO (GSE117498). Human bone marrow scATAC-seq data^40^ can be found at GEO (GSE96772).

## Code availability

CIBER toolkit is implemented in R and is available on Bioconductor and GitHub (https://github.com/Lan-lab/CIBER). Tutorials of CIBER usage cases are also available through GitHub (https://github.com/Lan-lab/CIBER/tutorials).

## Acknowledgements

The authors are grateful to Dr. Yeguang Chen from Tsinghua University for providing *Bcl11b*^flox/flox^ mice. We thank the High-Performance Computation platforms of Tsinghua University for providing computational resources.

## Author Contributions

L. X., J. W. and X. L. conceived the project. L. X. designed the models and theories. L. X. and H. X. implemented the code. L. X., H. X., N. Y., C. T. and S. Y. performed the analyses. J. W. designed the experiments and T. C. conducted the experiments. L. X. and T. C. generated the figures. M. Z., R. S., R. Y., W. Z. helped in designing the analyses and explaining the results. X. L. supervised the project. All authors contributed to the writing of the manuscript.

## Extended data

**Extended Data Fig. 1.**
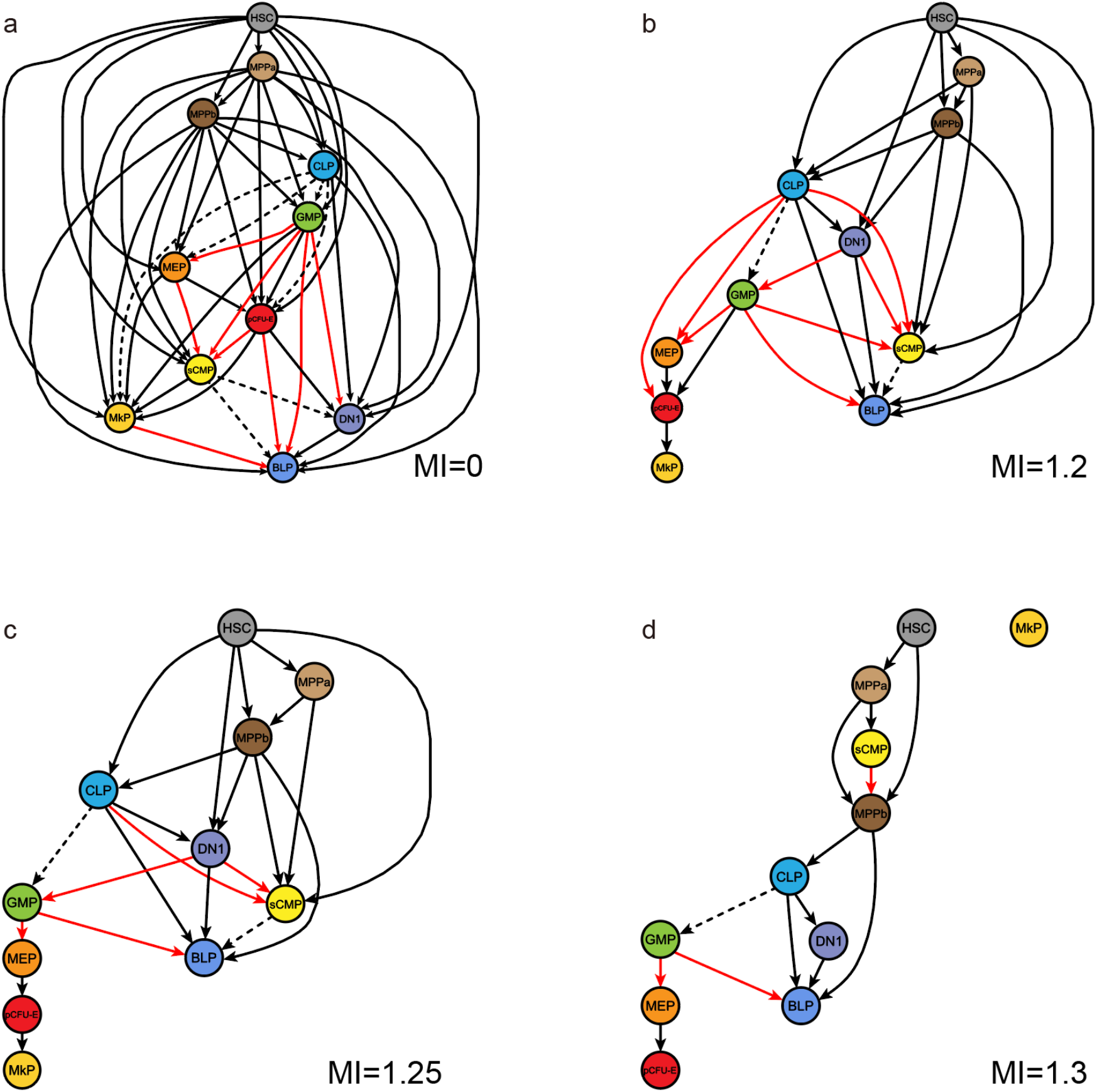
Typical Bayesian network structure learning algorithm is sensitive to parameters. **a-d**, Networks learned by the typical Bayesian network structure learning strategy MI-hc given different correlation thresholds with the same microarray dataset. The dashed edges represent possible unconventional interlineage branches, while the red edges represent less credible branches.

**Extended Data Fig. 2.**
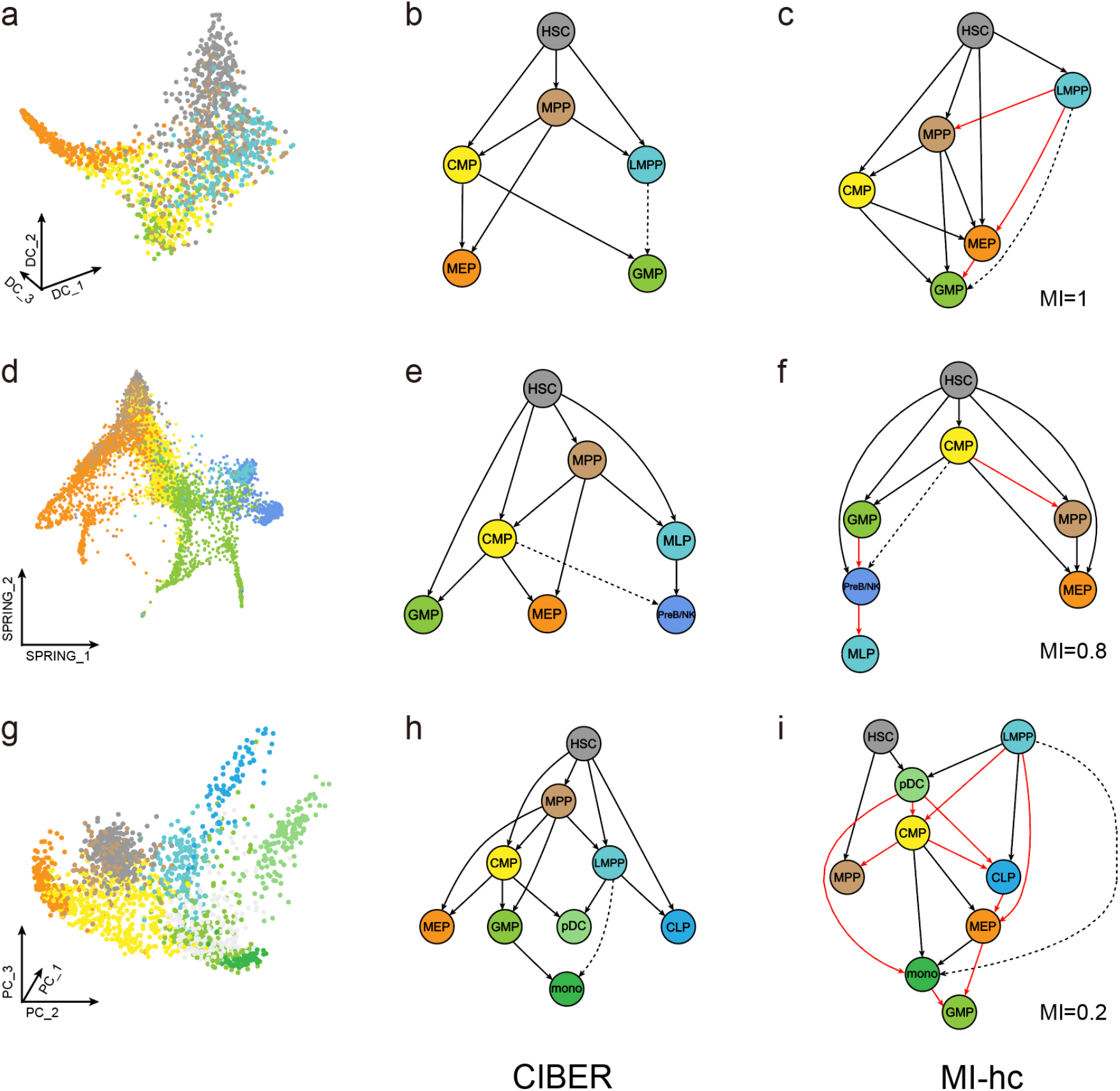
CIBER-learned structures conform to prior knowledge and reduction landscapes better than the MI-hc algorithm. Plots show low-dimensional landscapes^38–40^, CIBER and MI-hc results of (**a-c**) mouse scRNA-seq^38^, (**d-f**) human scRNA-seq^39^ and (**g-i**) human scATAC-seq^40^ datasets respectively. The dashed edges represent unconventional interlineage edges that were reported to be possible in previous studies, while the red edges represent less credible edges. The results show that compared to the typical Bayesian network structure learning algorithm MI-hc, the CIBER-learned results were consistent with conventional knowledge with a larger number and a higher percentage of edges and captured the bifurcation structure more clearly.

**Extended Data Fig. 3.**
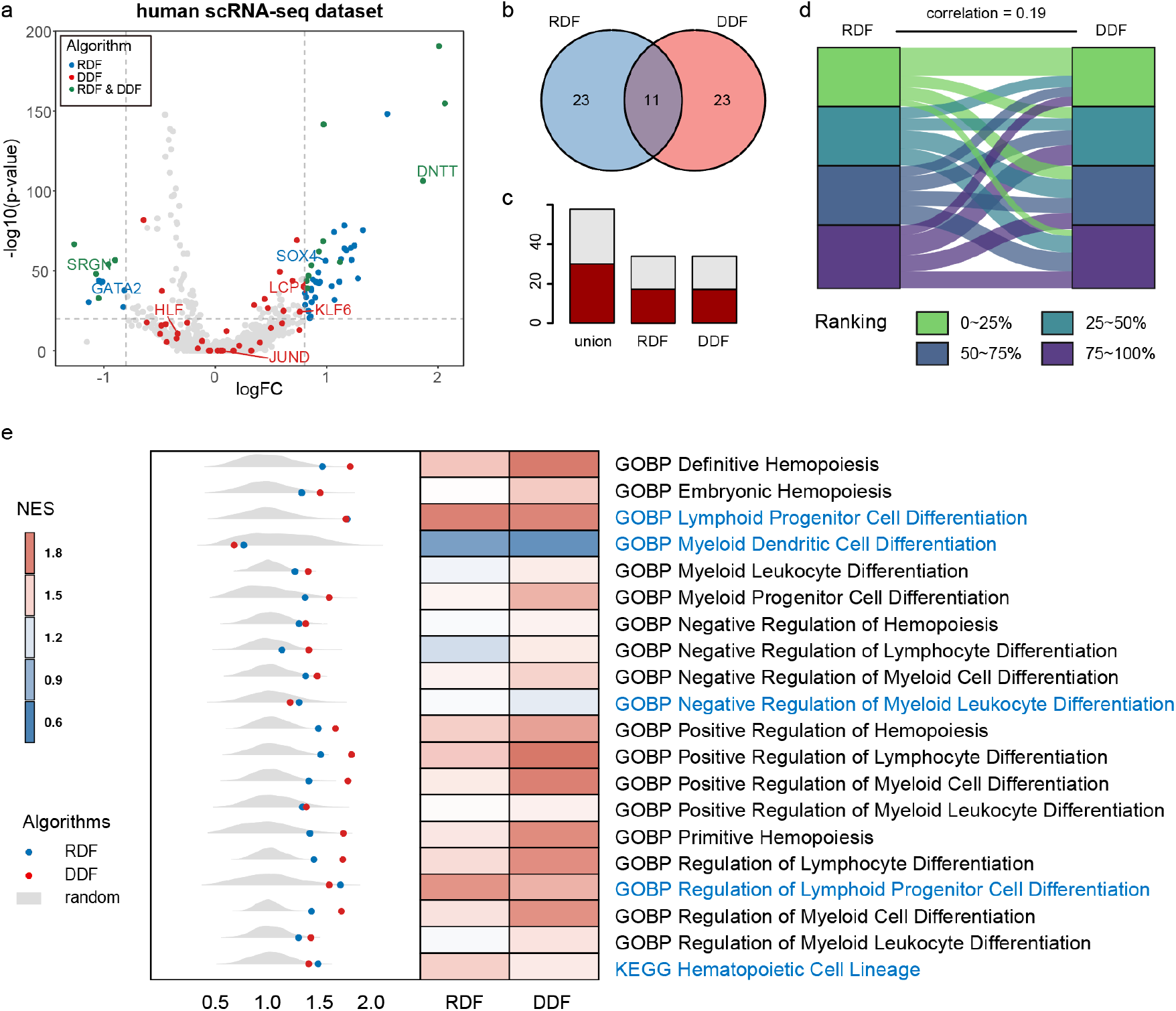
DDF identifies lineage driving features. The plots show the results on RDF and DDF gene lists obtained with the human scRNA-seq dataset. **a**, Features are allocated by log fold change and log p value obtained through RDF analysis; top RDF and DDF features are coloured. **b**, The number of shared features between the two feature sets. **c**, Literature search result shows that the DDF and RDF analyses identify different important features, in which the red parts represent the reported features. **d**, Correlation between the ranked lists. **e**, GSEA NES with terms related to haematopoiesis or haematopoietic differentiation in the GO and KEGG databases. The terms for which RDF list obtains the higher NES are coloured in blue. Dots in the left part represent the NES of RDF and DDF ranking lists, while the distributions show the background NES given randomly ranked gene lists.

**Extended Data Fig. 4.**
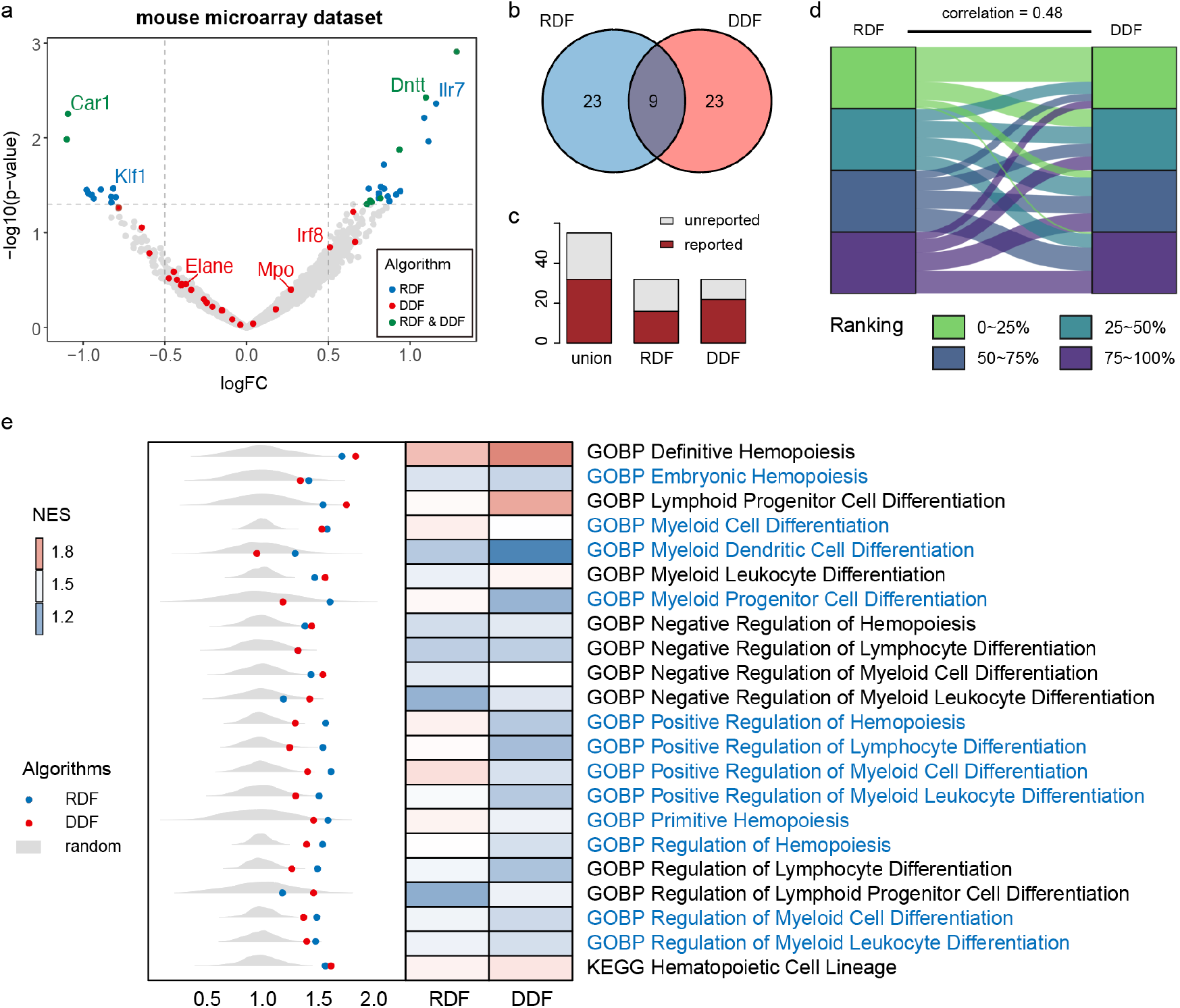
DDF results on the mouse microarray dataset. **a**, Features are allocated by log fold change and log p value obtained through RDF analysis; top RDF and DDF features are coloured. **b**, The number of shared top features between the two feature sets. **c**, Literature search result shows that the DDF and RDF analyses identify different important features, in which the red parts represent the reported features. **d**, Correlation between the ranked lists. **e**, GSEA NES with terms related to haematopoiesis or haematopoietic differentiation in the GO and KEGG databases. The terms for which RDF list obtains the higher NES are coloured in blue. Dots in the left part represent the NES of RDF and DDF ranking lists, while the distributions show the background NES given randomly ranked gene lists. The RDF list performed slightly better than the DDF list in 12 out of 22 terms. However, the literature search still showed that DDF analysis identified some important genes which are not significantly expressed, demonstrating that DDFs can complement RDFs to better identify genes crucial to cell differentiation.

**Extended Data Fig. 5.**
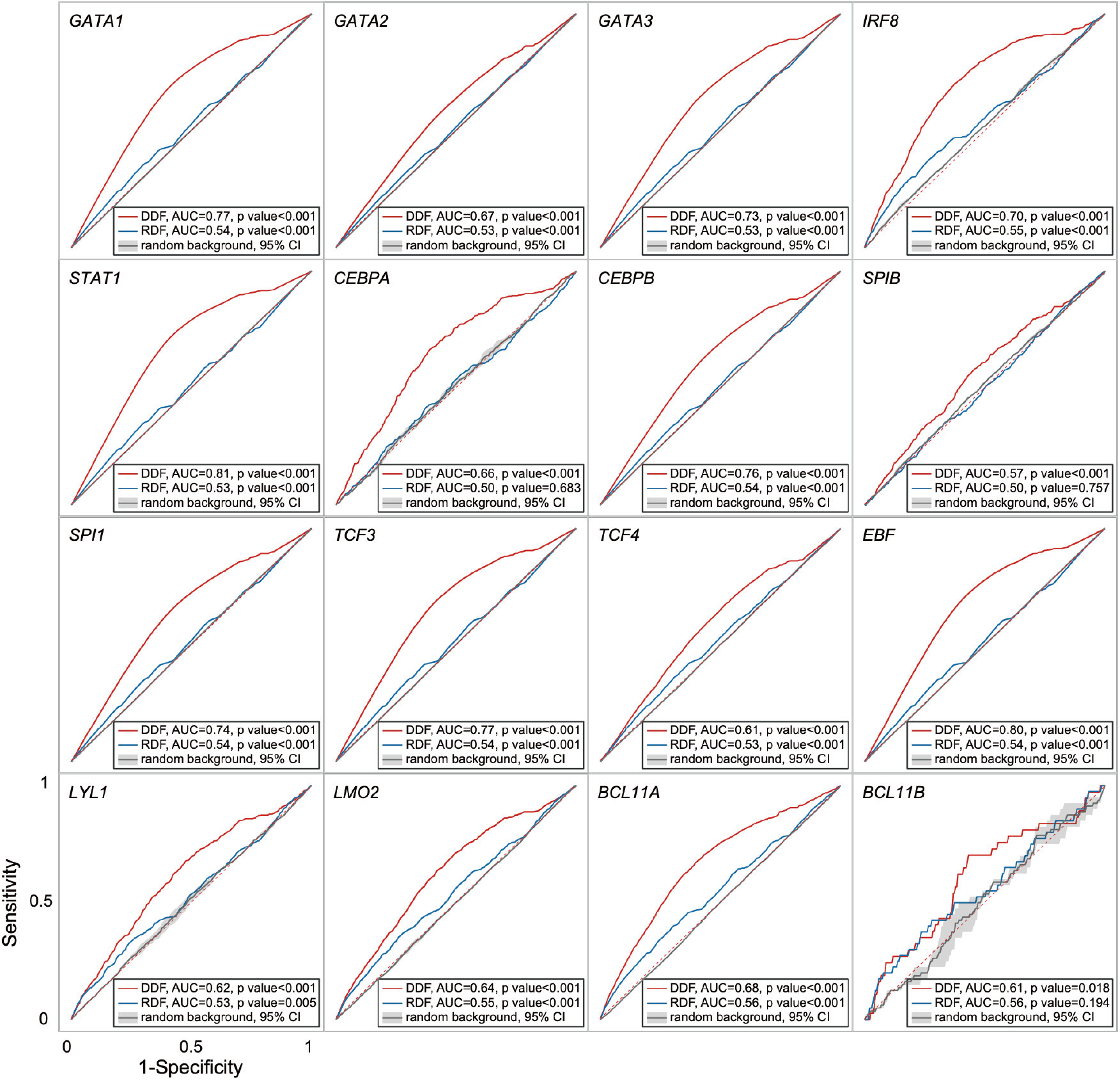
ROC curves on each gene set targeted by different TFs. Results show that DDF analysis ranks the genes targeted by the reported important TFs at higher positions compared to regular differential analysis, indicating that DDF analysis may provide insight into the potential mechanism of how important TFs impact the cellular differentiation process.

